# BH3 Profiling Identifies Selective BCL-2 Dependence of Adult Early T-cell Progenitor (ETP) Subtype of Acute Lymphoblastic Leukemia (ALL) Patients

**DOI:** 10.1101/2021.08.19.456932

**Authors:** Elyse A. Olesinski, Karanpreet Singh Bhatia, Jacqueline Garcia, Nitin Jain, Anthony Letai, Marina Konopleva, Shruti Bhatt

**Author notes:** Co-corresponding **Corresponding Authors**: Shruti Bhatt, PhD, Address: Department of Pharmacy, Science Drive 3, Block S7 Room 02-16, Singapore 129788, Marina Konopleva, MD, PhD, Department of Leukemia, University of Texas M.D. Anderson Cancer Center, Houston, Texas, United States of America, 1400 Holcombe Blvd., FC3.3048.

## Abstract

Early T-cell precursor (ETP) acute lymphoblastic leukemia (T-ALL) is a high-risk subtype of T-ALL and is associated with poor survival outcomes with chemotherapy. We previously showed that maturation stage of thymocytes distinguishes pro-survival dependencies of pediatric ETP-ALL (BCL-2 dependent) from non-ETP-ALL (BCL-XL dependent). Comparable data in adults are lacking. Our present study focuses on characterizing functional dependencies of adult ETP-ALL and T-ALL on BCL-2 family proteins. Using BH3 profiling on primary tumors, we report that similar to pediatric ALL, adult ETP-ALL is primarily dependent on BCL-2; however, unlike pediatric ALL, adult non-ETP-ALL is co-dependent on both BCL-2 and BCL-XL for survival. By measuring direct mitochondrial permeabilization and cell viability assays, we validated that BH3 profiling predicted on-target cytotoxicity of venetoclax in adult ETP-ALL and navitoclax plus venetoclax in adult T-ALL. These findings provide pre-clinical evidence for venetoclax and navitoclax as potentially efficacious combination therapy for adults with T-ALL.

**Statement of Significance:** Adults with relapsed T-ALL have poor prognosis and represent an unmet need for novel treatments. We report that adult ETP-ALL is dependent on BCL-2 for survival, similar to pediatric ETP-ALL. However, adult non-ETP-ALL is heterogeneously dependent on both BCL-2 and BCL-XL. We characterize key dependencies on BCL-2 family proteins to direct BH3 mimetics therapy in T-ALL patients.

## Introduction

T-cell acute lymphoblastic leukemia (ALL) results from malignant transformation of immature T-cells, accounting for 10-15% of childhood and 20-25% of adult ALL cases.^1^ Although long-term survival rates for standard risk childhood T-cell ALL (T-ALL) have shown striking improvements to 90%, response outcomes in adults remain much lower, possibly due to high induction therapy-related toxicities.^2^ In general, the causes for poor response in adult T-ALL are incompletely understood.

Among T-ALL, a high-risk subtype originating from clonal expansion of recently immigrated thymocytes with early differentiation arrest, defined as early T-cell precursor (ETP) leukemias, has a significantly worse outcome in adults.^3^ ETP-ALL retains multipotent differentiation capacities and resemblance to hematopoietic stem cells or myeloid progenitor cells. Targeted next-generation sequencing^4^ showed that adult ETP-ALL presents a similar mutation profile as its pediatric equivalent. Recent data showed that, similar to children, adult ETP-ALL has inferior outcomes to chemotherapy than non-ETP-ALL. However, implementation of allogenic transplant after achieving first complete response with standard chemotherapy regimens improved the overall prognosis of patients with ETP-ALL, similar to that of the rest of the T-ALL cohort in high-risk patients.^4^ However, the 5-year overall survival rate of adults with ETP-ALL is only 49%.^4^ These outcomes highlight the unmet need for new therapeutic approaches for adult ETP-ALL.

BH3 mimetics, small molecule antagonists of BCL-2 family proteins, have recently shown clinical success in various hematological malignancies, including chronic lymphocytic leukemia (CLL), acute myeloid leukemia (AML), and ALL. Studies from our group^5–8^ and several others^9–11^ showed that BH3 profiling can predict tumor cells’ functional dependency on BCL-2. In turn, BH3 profiling identified BCL-2 dependence directed clinical success of the FDA-approved venetoclax (BCL-2 inhibitor) in CLL^12^ and AML.^6,7^ BH3 profiling method measures functional interactions between BCL-2 family proteins by using BH3 peptides that mimic BH3 domains present in pro-apoptotic proteins.^13^ In practice, BH3 profiling involves exposing the mitochondria to synthetic BH3 peptides and subsequently measuring the ensuing mitochondrial outer membrane permeabilization (MOMP). When selective peptides are used, e.g. BAD, heightened mitochondrial sensitivity indicates greater dependence on an individual anti-apoptotic protein, e.g. BCL-2 and BCL-XL.

In a previous study from our group, we reported that pediatric ETP-ALL is BCL-2 dependent, while pediatric T-ALL is BCL-XL dependent.^14^ Since therapeutic success of adult ALL is different from pediatric ALL, the aim of this study was to identify anti-apoptotic BCL-2 family protein dependencies and sensitivities to BH3 mimetics of adult ETP-ALL and T-ALL lymphoblasts using BH3 profiling.

## Results

### BH3 profiling identifies BCL-2 dependence in adult ETP-ALL cells and sensitivity to venetoclax and navitoclax

To determine the anti-apoptotic dependencies of adult ETP-ALL, we performed BH3 profiling on 13 primary adult ETP-ALL samples obtained at diagnosis (Figure 1A; Table 1). We exploited the different binding affinities of BAD BH3 and HRK BH3 peptides to distinguish between BCL-2 dependence (high BAD, low/no HRK signaling) and BCL-XL dependence (equal BAD and HRK signaling) (Figure 1B). Since the BAD peptide interacts with BCL-2, BCL-XL, and BCL-W, and the HRK peptide interacts with BCL-XL alone, a rough index for selective BCL-2 dependence can be obtained by subtracting the response to the HRK peptide from the response to the BAD peptide (BAD-HRK). All 13 samples showed robust mitochondrial depolarization in response to the BAD peptide as compared to the HRK peptide and DMSO control, suggesting primary dependence on pro-survival BCL-2 (Figure 1C). Despite heterogeneity in mitochondrial priming responses among samples, the mean of BAD-induced cytochrome c release caused by all samples was significantly higher compared to the HRK peptide (p<0.0001, Figure 1C). We found no association between BAD and HRK (Spearman r=0.071, p=0.82), but BAD and BAD-HRK had a statistically significant association (Spearman r=0.57, p<0.05; Figure 1D). This suggests that mitochondrial depolarization caused by the BAD peptide corresponds to more BCL-2 dependence than BCL-XL dependence. To further validate BCL-2 dependence of ETP blasts, we measured direct mitochondrial sensitivity to the BH3 mimetics venetoclax (which selectively antagonizes BCL-2) and navitoclax (which antagonizes BCL-2 and BCL-XL). Since both drugs antagonize BCL-2, both drugs showed increased cytochrome c release as compared to DMSO, using BH3 profiling assays. We found venetoclax to be a better inducer of mitochondrial priming compared to navitoclax, which can be explained by the higher binding affinity of venetoclax (K_i_<0.010 nM)^15^ over navitoclax (K_i_=0.044 nM)^15^ for BCL-2 (p<0.001; Figure 1E).

**Table 1.**
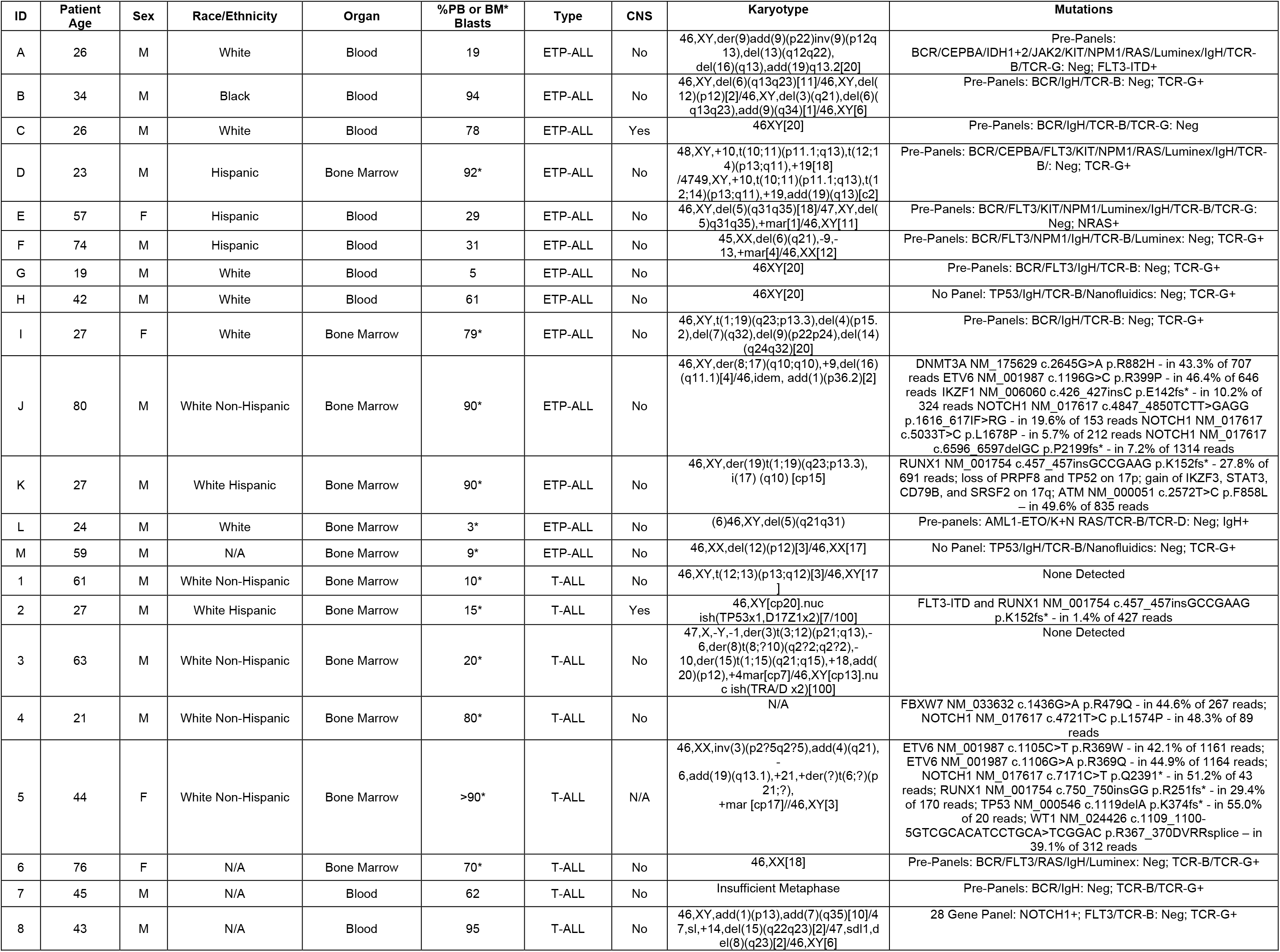
Clinical patient characteristics and ALL subtypes. Samples from Dana-Farber Cancer Institute were screened using a CLIA-approved next-generation sequencing assay (BWH Rapid Heme Panel) based on the TruSeq Custom Amplicon kit from Illumina.^20^ This method assessed 95 genes with hot spot codons for oncogenes and full coding regions for tumor suppressor genes. Samples from MD Anderson were screened following a specific pre-panel to determine karyotype and mutations. No gene panel was used for sample M, and sample 8 was screened using a 28 gene panel. * percent blasts in bone marrow versus peripheral blood. Abbreviations: T-ALL, typical ALL; ETP-ALL, early T-cell progenitor ALL.

**Figure 1.**
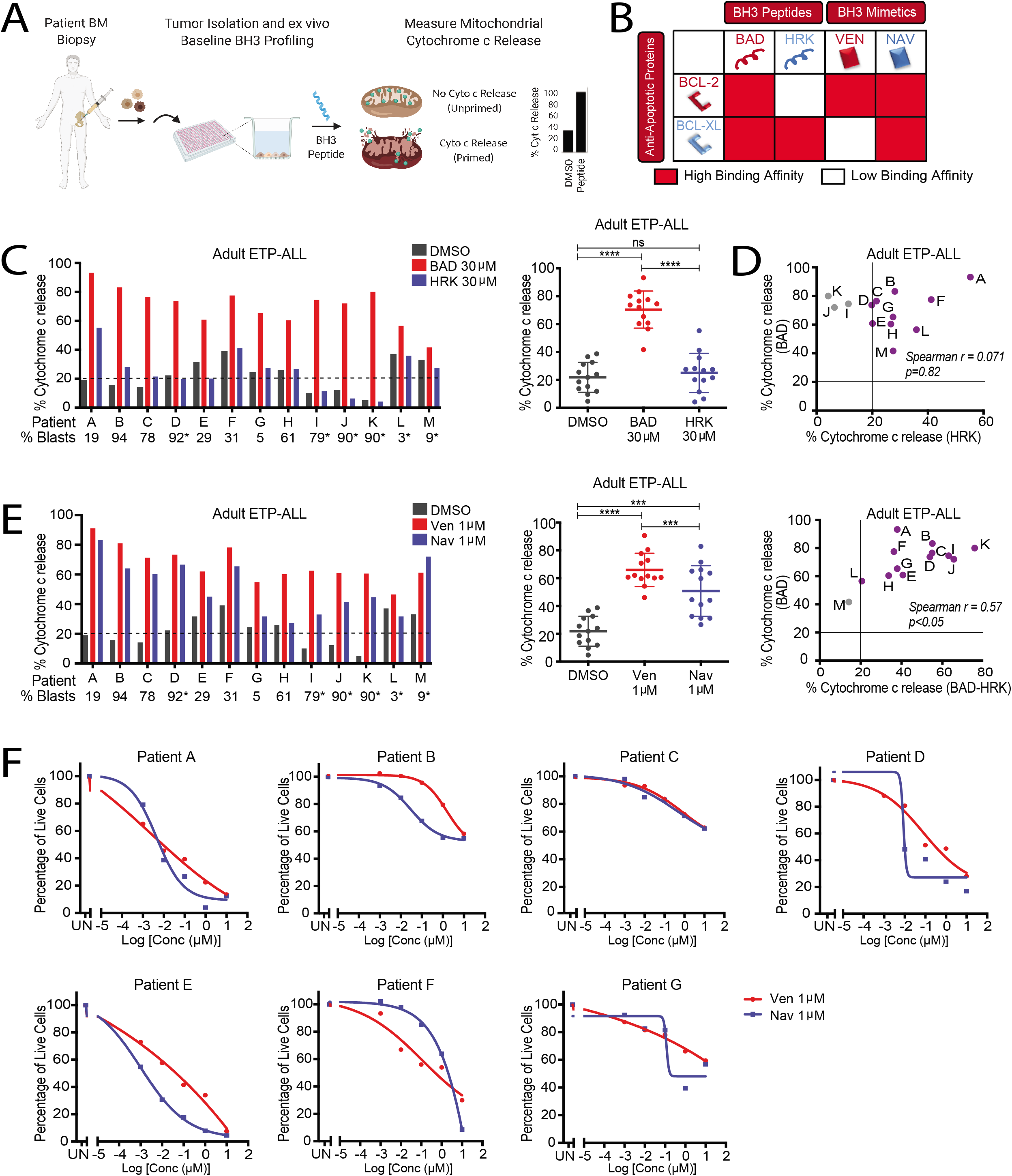
Adult ETP-ALL have increased dependency on BCL-2 rather than BCL-XL for survival. **A**. Schematic for baseline BH3 profiling. **B**. Binding affinities of BH3 peptides (BAD and HRK) and BH3 mimetics (venetoclax and navitoclax) for anti-apoptotic proteins BCL-2 and BCL-XL. Red, high-affinity binding; white, low-affinity binding. **C**. BH3 profiles of adult ETP-ALL primary samples in response to BAD and HRK peptides. One-way ANOVA analysis demonstrates a statistical difference between the percent cytochrome c release between BAD versus DMSO (p<0.0001) and BAD versus HRK (p<0.0001). There is no statistical difference between HRK versus DMSO (p=0.3480). * percent blasts in bone marrow versus peripheral blood. The dotted line represents threshold (20%) for significant priming. **D**. Spearman correlation between the percent cytochrome c release for BAD versus HRK (r=0.071, p=0.82) and between BAD versus BAD-HRK (r=0.57, p<0.05). **E**. BH3 profiles of adult ETP-ALL primary samples in response to venetoclax and navitoclax. One-way ANOVA analysis demonstrates a statistical difference between the percent cytochrome c release between venetoclax versus DMSO (p<0.0001), navitoclax versus DMSO (p<0.001), and venetoclax versus navitoclax (p<0.001) in adult ETP-ALL primary patient samples. **F**. Cell viability analysis for adult ETP-ALL primary samples treated with increasing concentrations of venetoclax and navitoclax for 8 hours. Data are plotted as the percentage of luminescence compared to DMSO controls. UN = untreated.

Our results from BH3 profiling showing primary dependence of adult ETP-ALL on BCL-2 led to the hypothesize that primary ETP-ALL tumors are sensitive to BH3 mimetics inhibiting BCL-2 protein. We performed cell viability assays by exposing ETP-ALL primary tumors to short-term treatment with venetoclax and navitoclax. We specifically focused on venetoclax and navitoclax as BH3 mimetics drugs because of their clinical relevance and clinical advancement in hematological cancer treatment. Indeed, while response was heterogeneous, all samples tested demonstrated sensitivity to both venetoclax and navitoclax (Figure 1F). These findings demonstrate that consistent with BCL-2 dependence, adult ETP-ALL can be efficiently targeted for apoptosis via venetoclax.

### BH3 profiling reveals BCL-2/BCL-XL co-dependence in adult T-ALL and sensitivity to venetoclax and navitoclax

Having shown selective apoptotic dependence of adult ETP-ALL on BCL-2, we next evaluated whether adult non-ETP T-ALL also shows survival dependency on specific BCL-2 family proteins. We performed BH3 profiling on 8 primary adult T-ALL samples collected at diagnosis (Table 1) using identical BH3 peptide panels and concentrations. Contrary to adult ETP-ALL, 7 out of 8 samples showed mitochondrial depolarization in response to both BAD and HRK peptides compared to DMSO controls (p<0.05, Figure 2A).

**Figure 2.**
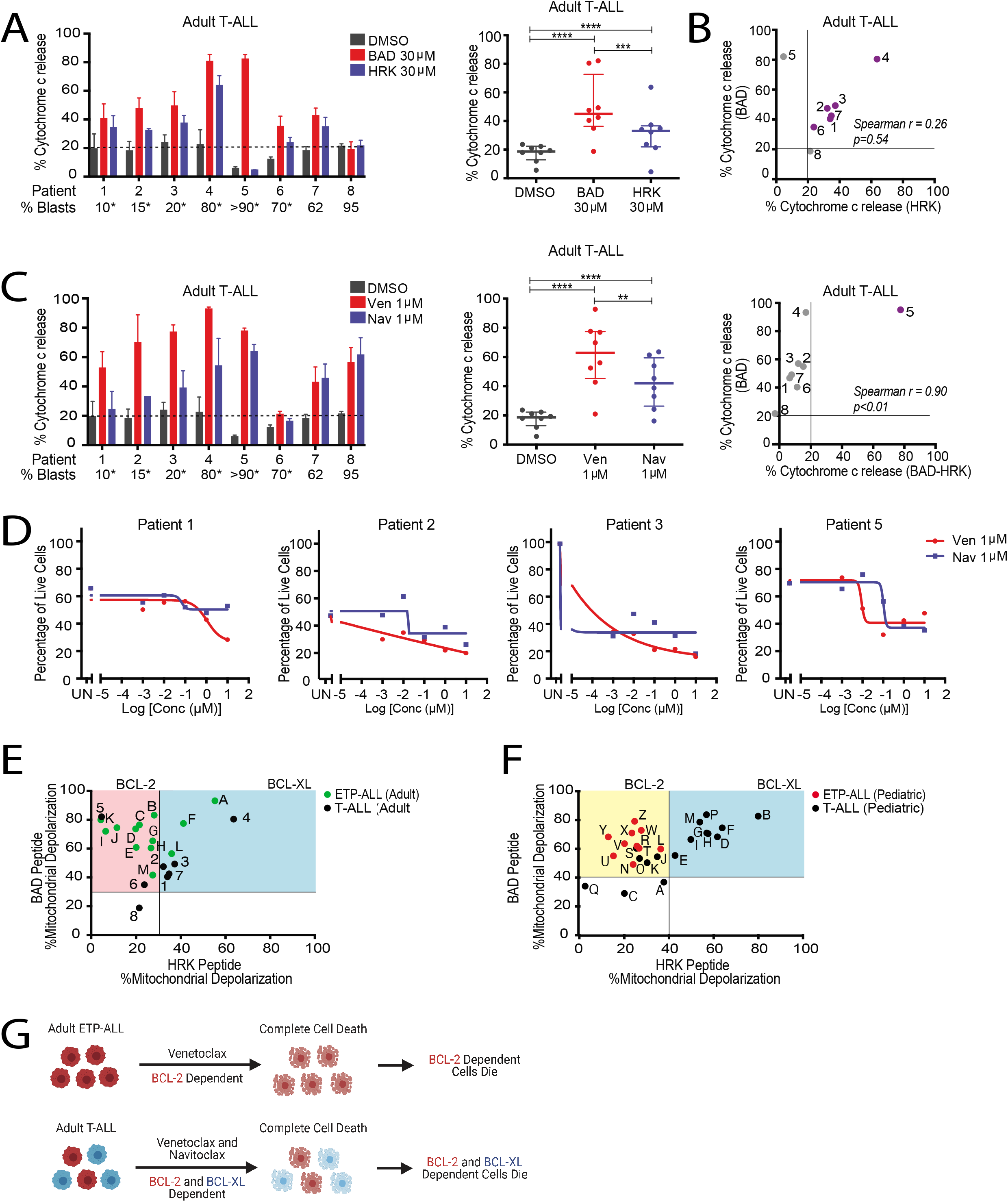
Adult T-ALL specimens are dependent on both BCL-2 and BCL-XL for survival. **A**. BAD versus HRK BH3 profiles of adult T-ALL primary samples. Two-way ANOVA analysis demonstrates a statistical difference between the percent cytochrome c release between BAD versus DMSO (p<0.01) and HRK versus DMSO (p<0.01). The mean ± SD of three independent experiments before treatment are graphed. * percent blasts in bone marrow versus peripheral blood. The dotted line represents the threshold for significant priming determined by DMSO±3xSD. **B**. Spearman correlation between the percent cytochrome c release for BAD versus HRK (r=0.26, p=0.54) and between BAD versus BAD-HRK (r=0.90, p<0.01). **C**. Venetoclax versus navitoclax BH3 profiles of adult T-ALL primary samples. Two-way ANOVA analysis demonstrates a statistical difference between the percent cytochrome c release between venetoclax versus DMSO (p<0.0001), navitoclax versus DMSO (p<0.0001), and venetoclax versus navitoclax (p<0.01). The mean ± SD of three independent experiments are graphed. **D**. Cell viability analysis for adult T-ALL primary samples treated with increasing concentrations of venetoclax and navitoclax for 8 hours. Data are plotted as the percentage of luminescence compared to DMSO controls. UN = untreated. **E**. A dot plot of the BAD peptide response versus the HRK peptide response in adult ETP-ALL (green) and T-ALL (black). Red shows probable BCL-2 dependence, and blue shows probable BCL-XL dependence. **F**. Adapted from Ni Chongaile et al. 2015: Dot plot of the BAD peptide response versus the HRK peptide response in pediatric ETP-ALL (red) and T-ALL (black). Yellow shows probable BCL-2 dependence, and blue shows probable BCL-XL dependence. **G**. Proposed model schematic for targeting of ETP-ALL and T-ALL based on BCL-2 and BCL-XL protein dependencies.

Since T-ALL primary tumors showed induction in mitochondrial priming to both HRK and BAD peptides, it is likely that T-ALL blasts may be co-dependent on BCL-2 and BCL-XL. We next performed a comparison between mitochondrial depolarization response to BAD vs BAD-HRK or HRK alone to determine whether mitochondrial priming response to BAD is driven by BCL-2 or BCL-XL or both. Similar to ETP-ALL, we found poor association between BAD and HRK (Spearman r=0.26, p=0.54), but BAD and BAD-HRK showed a significant association (Spearman r=0.90, p<0.01; Figure 2B), indicating BCL-2 dependence. In addition, simultaneous enhancement in the HRK peptide sensitivity compared to DMSO indicates that adult T-ALL are co-dependent on BCL-2 and BCL-XL. We next measured direct mitochondrial priming sensitivity to venetoclax and navitoclax using BH3 profiling assay. Both venetoclax and navitoclax caused similar cytochrome c release compared to DMSO, indicating that T-ALL tumors are dependent on BCL-2 and BCL-XL for survival (Figure 2C). Similar to ETP-ALL, venetoclax elicited slightly higher priming across samples than navitoclax (p<0.01; Figure 2C).

Because we observed robust mitochondrial depolarization in responses to both BAD and HRK peptides, we hypothesized that T-ALL tumors would show enhanced cytotoxic response to BH3 mimetics drugs targeting BCL-2 and BCL-XL. We performed cell viability assays by exposing T-ALL primary tumors to short-term treatment with venetoclax and navitoclax. Of note, only 50% of primary T-ALL tumors maintained >50% baseline viability that precluded us from performing cytotoxicity experiments on all tumors. Nonetheless, we found that the 4 samples that met analyzable cut-off criteria (>50% baseline viability) demonstrated sensitivity to venetoclax and navitoclax, which reiterates that antagonism of BCL-2 and BCL-XL anti-apoptotic proteins may improve therapeutic outcomes (Figure 2D).

Finally, we asked if there is a difference in dependencies between adult ETP-ALL and T-ALL. Hence, we compared mitochondrial depolarization profiles of both ETP-ALL and T-ALL tumors in response to BAD and HRK peptides (Figure 2E). In ETP-ALL, cytochrome c release was caused by the BAD peptide alone, while in T-ALL, both BAD and HRK peptides induced cytochrome c release (Figure 2E). This suggests that ETP-ALL primary tumors are primarily BCL-2 dependent, while T-ALL primary tumors are co-dependent on BCL-2 and BCL-XL.

Since we previously carried out similar study primarily on pediatric ALL cohort,^14^ we compared both cohorts to ask if there was age-dependent difference in mitochondrial priming between adult vs pediatric ALL. We found that mean of BAD-induced mitochondrial priming in pediatric ALL cohort exceeded adult T-ALL counterparts (70.4% vs 49.5%; p,0.05; Figure 2E and 2F). However, mitochondrial sensitivity to BAD peptide was not significantly different between pediatric vs adult in ETP-ALL tumors. Collectively, pediatric ETP-ALL was most sensitive to BAD-induced mitochondrial depolarization compared to T-ALL (Figures 2E and 2F).

## Discussion

Pediatric T-ALL has better long-term survival outcomes than adult (80-90% vs. 40%).^2^ BH3 mimetics, small molecule antagonists of anti-apoptotic BCL-2 family proteins, have shown remarkable clinical success in the treatment of hematological malignancies. In particular, BCL-2 antagonist venetoclax received FDA approval for treatment of AML and CLL. We previously showed that pediatric ETP-ALL, an ALL subtype with poor prognosis, is dependent on BCL-2, while typical T-ALL is dependent on BCL-XL.^14^ These findings importantly led to the clinical trial of venetoclax in combination with hyperCVAD in relapsed/refractory (R/R) T-ALL patients.^12,14,16,17^ However, whether adult patients with T-ALL also show the selective pattern of anti-apoptotic dependence related to differentiation stage of T-cell has not been studied. Therefore, we investigated the survival dependencies of adult ETP-ALL and T-ALL on BCL-2 family proteins as well as identified sensitivities to different BH3 mimetics using BH3 profiling.

We found that adult ETP-ALL is more dependent on BCL-2 than on BCL-XL for survival. This finding is consistent with our previous study carried out in pediatric ETP-ALL. However, unlike pediatric T-ALL, which showed selective dependence on BCL-XL, adult T-ALL was found to be co-dependent on BCL-2 and BCL-XL. Despite differences in selectivity of dependence, these findings importantly emphasize that adult T-ALL cells may be sensitive to dual inhibition of BCL-2 and BCL-XL, broadening the range of potentially effective agents. With these cellular dependencies in mind, we propose that adult ETP-ALL may be sensitive to treatment with venetoclax, while adult T-ALL may be sensitive to combination of venetoclax and navitoclax (Figure 2F). Noticeably, comparison of ETP- and T-ALL samples demonstrated that, irrespective of BCL-2 or BCL-XL dependence, the degree of mitochondrial priming was lower in T-ALL as compared to ETP-ALL samples (Figure 2E).

While our study was focused on directing the use of BH3 mimetics in the upfront setting, the results from a recent Phase I study (NCT03181126) in R/R B-ALL and T-ALL using a combination of venetoclax and navitoclax showed that dual dependence on BCL-2 and BCL-XL is maintained in the relapse settings.^18^ The reported overall response rate in R/R ALL within all subgroups was 59.9% (n=28/47), with pediatric subgroups showing a higher overall response rate of 75% (9/12) on combination therapy with venetoclax and navitoclax.^18^ Although data is pending on whether combination therapy is superior to venetoclax alone, our findings demonstrate, in concordance with ongoing research, that targeting dual dependency on BCL-2/BCL-XL anti-apoptotic proteins using BH3 mimetics improve outcomes of adult T-ALL.

Despite successfully characterizing the functional vulnarabilties of adult ETP-ALL and T-ALL with BH3 profiling and, our study was limited to small sample size due to the rarity of occurrence. Nonetheless, BH3 profiling distinctly identified BCL-2 and BCL-XL dependencies that correlated with BH3 mimetics sensitivity. Also, compared to frank cell death or cell viability measurements requiring longer incubation times, rapid measurement in mitochondrial depolarization assessments via BH3 profiling offered the major advantage of descriptive data despite low cell recovery. In fact, out of our 21 primary adult ETP-ALL and T-ALL tumors, 10 failed cytotoxicity readouts due to poor viability at baseline after 8 hours of *ex vivo* culture. As expected, we observed intra-patient variability in mitochondrial priming responses to BH3 peptides and mimetics. Given the heterogeneous nature of blood cancers and rarity of ALL in adults, we prioritized looking into BCL-2 and BCL-XL dependence via BH3 profiling in our small sample size. However, other unknown interactions may influence the efficacy of BCL-2 and BCL-XL inhibitors in this patient population, such as MCL-1, BFL-1, and BMF. Moving forward, BH3 profiling will continue to be a valuable biomarker functional tool for identifying protein dependencies and drug vulnerabilities in adult ALL patients.

## Acknowledgments

The authors thank all the patient and families that consented to provide samples for this study at Dana-Farber Cancer Institute and MD Anderson Cancer Center. SB is a recipient of a Career Development Award from the Leukemia and Lymphoma Society and Basic Cancer Research Fellowship award from AACR. MK was supported by R01CA241191. JSG was supported by the National Cancer Institute of the National Institutes of Health under Award Number K08CA245209 and NIH/NCI SPORE in Myeloid Malignancies Grant Number 1P50CA206963. The content is solely the responsibility of the authors and does not necessarily represent the official views of the NIH or NCI.

## Authorship Contributions

SB, MK and AL designed the laboratory studies. JSG and NJ coordinated the study, provided clinical samples, regulatory oversight, and collected the study data. SB and EAO performed BH3 profiling and cell death assays, and wrote the manuscript. SB, EAO and KSB analyzed the data. All authors read, critically reviewed, and approved the final manuscript.

## Methods

### Human Subjects

ALL patient clinical characteristics and subtype are described in detail in Table 1. All samples were obtained after informed patient consent under IRB approved Dana-Farber Cancer Institute and MD Anderson Cancer Center collection protocols. All samples were obtained at the diagnosis stage. ETP-ALL was immunophenotypically characterized as CD1a negative (<5%), CD8 negative (<5%), absent or weak CD5 (<75%), with the presence of at least 1 myeloid or stem cell marker (>25%), such as CD117, CD34, HLA-DR, CD13, CD33, CD11b, or CD65.

### Intracellular BH3 (iBH3) profiling

Peripheral blood samples or bone marrow aspirates were collected from patients with ETP-ALL or T-ALL. Mononuclear cells were obtained using a Ficoll-pacque Plus (GE Health Care) and viably frozen before subjecting to iBH3 profiling, as described earlier.^5^ Cells were counted to determine recovery using trypan blue positivity stain. Samples with >70% viability were considered suitable for further processing. Cell staining with 1:100 live/dead zombie yellow was first carried out (Biolegend) in PBS at room temperature for 15 minutes. After incubation, cells were washed with PBS and stained with 1:100 CD45 (BD Biosciences) in FACS buffer (2% FBS in PBS) on ice for 30 minutes. BH3 profiling was performed.^19^ Synthetic BH3 peptides were combined with MEB buffer (150 mM mannitol, 5 mM succinate, 50 mM KCI, 0.02 mM EGTA, 0.02 mM EDTA, 10 mM HEPES-KOH pH 7.5, and 0.1% BSA) and 0.001% digitonin. Cells were incubated with peptides for 60 minutes to facilitate plasma membrane permeabilization. Subsequently, cells were fixed with 4% formaldehyde at room temperature for 15 minutes, followed by neutralization at room temperature for 10 minutes with N2 buffer (1.7 M Tris, 1.25 M Glycine pH 9.1). Finally, cells were incubated overnight with anti-cytochrome c Alexafluor 647 antibody (Clone 6H2.B4, Biolegend) in staining buffer (2% Tween20, 10% BSA, PBS). To measure cytochrome c loss due to BH3 peptide sensitivity, FACS analysis was performed by gating on DMSO negative controls to describe 100% cytochrome c retention. Alamethicin was used as a positive control for 100% cytochrome c release. Blasts were confirmed by CD45 dim/SSC-low. The following equation was used to calculate cytochrome c loss: (100 – percent of cells within cytochrome c retention gate).

### Cell Viability Assays

ETP-ALL and T-ALL cells in RPMI media were plated in 96-well plates and treated with venetoclax or navitoclax at concentrations ranging from 0.000001 μM to 10 μM. Cells were incubated with the drugs at 37°C for 8 hours, after which cell viability was measured using CellTiter-Glo reagents (Promega) following the manufacturer’s protocol. Absolute viability values were converted to percentage viability versus DMSO control or versus treatment in a non-linear fit of Log (inhibitor) versus response. GraphPad Prism version 9.0 was used to perform variable slope (four parameters) and to obtain IC50 metrics.

## References

1. Van Vlierberghe P, Ferrando A. The molecular basis of T cell acute lymphoblastic leukemia. J Clin Invest. 2012;122(10):3398–3406. doi:10.1172/JCI61269

2. Inaba H, Greaves M, Mullighan CG. Acute lymphoblastic leukaemia. Lancet. 2013;381(9881). doi:10.1016/S0140-6736(12)62187-4

3. Jain N, Lamb AV, O’Brien S, et al. Early T-cell precursor acute lymphoblastic leukemia/lymphoma (ETP-ALL/LBL) in adolescents and adults: a high-risk subtype. Blood. 2016;127(15):1863–1869. doi:10.1182/blood-2015-08-661702

4. Bond J, Graux C, Lhermitte L, et al. Early Response-Based Therapy Stratification Improves Survival in Adult Early Thymic Precursor Acute Lymphoblastic Leukemia: A Group for Research on Adult Acute Lymphoblastic Leukemia Study. J Clin Oncol Off J Am Soc Clin Oncol. 2017;35(23):2683–2691. doi:10.1200/JCO.2016.71.8585

5. Bhatt S, Pioso MS, Olesinski EA, et al. Reduced Mitochondrial Apoptotic Priming Drives Resistance to BH3 Mimetics in Acute Myeloid Leukemia. Cancer Cell. 2020;38(6):872–890.e6. doi:10.1016/j.ccell.2020.10.010

6. Pan R, Hogdal LJ, Benito JM, et al. Selective BCL-2 inhibition by ABT-199 causes on-target cell death in acute myeloid leukemia. Cancer Discov. 2014;4(3):362–375. doi:10.1158/2159-8290.CD-13-0609

7. Konopleva M, Pollyea DA, Potluri J, et al. Efficacy and Biological Correlates of Response in a Phase II Study of Venetoclax Monotherapy in Patients with Acute Myelogenous Leukemia. Cancer Discov. 2016;6(10):1106–1117. doi:10.1158/2159-8290.CD-16-0313

8. Anderson MA, Deng J, Seymour JF, et al. The BCL2 selective inhibitor venetoclax induces rapid onset apoptosis of CLL cells in patients via a TP53-independent mechanism. Blood. 2016;127(25):3215–3224. doi:10.1182/blood-2016-01-688796

9. Gilmore A, King L. Emerging approaches to target mitochondrial apoptosis in cancer cells. F1000Research. 2019;8. doi:10.12688/f1000research.18872.1

10. Matulis SM, Boise LH. BCL2 dependency in diffuse large B-cell lymphoma: it’s a family affair. Haematologica. 2020;105(8):1993–1996. doi:10.3324/haematol.2020.253591

11. Kapoor I, Bodo J, Hill BT, Hsi ED, Almasan A. Targeting BCL-2 in B-cell malignancies and overcoming therapeutic resistance. Cell Death Dis. 2020;11(11):1–11. doi:10.1038/s41419-020-03144-y

12. Del Gaizo Moore V, Schlis KD, Sallan SE, Armstrong SA, Letai A. BCL-2 dependence and ABT-737 sensitivity in acute lymphoblastic leukemia. Blood. 2008;111(4):2300–2309. doi:10.1182/blood-2007-06-098012

13. Certo M, Del Gaizo Moore V, Nishino M, et al. Mitochondria primed by death signals determine cellular addiction to anti-apoptotic BCL-2 family members. Cancer Cell. 2006;9(5):351–365. doi:10.1016/j.ccr.2006.03.027

14. Chonghaile TN, Roderick JE, Glenfield C, et al. Maturation stage of T-cell acute lymphoblastic leukemia determines BCL-2 versus BCL-XL dependence and sensitivity to ABT-199. Cancer Discov. 2014;4(9):1074–1087. doi:10.1158/2159-8290.CD-14-0353

15. Souers AJ, Leverson JD, Boghaert ER, et al. ABT-199, a potent and selective BCL-2 inhibitor, achieves antitumor activity while sparing platelets. Nat Med. 2013;19(2):202–208. doi:10.1038/nm.3048

16. Khaw SL, Suryani S, Evans K, et al. Venetoclax responses of pediatric ALL xenografts reveal sensitivity of MLL-rearranged leukemia. Blood. 2016;128(10):1382–1395. doi:10.1182/blood-2016-03-707414

17. Alford SE, Kothari A, Loeff FC, et al. BH3 Inhibitor Sensitivity and Bcl-2 Dependence in Primary Acute Lymphoblastic Leukemia Cells. Cancer Res. 2015;75(7):1366–1375. doi:10.1158/0008-5472.CAN-14-1849

18. Pullarkat VA, Lacayo NJ, Jabbour E, et al. Venetoclax and Navitoclax in Combination with Chemotherapy in Patients with Relapsed or Refractory Acute Lymphoblastic Leukemia and Lymphoblastic Lymphoma. Cancer Discov. Published online January 1, 2021. doi:10.1158/2159-8290.CD-20-1465

19. Ryan J, Montero J, Rocco J, Letai A. iBH3: simple, fixable BH3 profiling to determine apoptotic priming in primary tissue by flow cytometry. Biol Chem. 2016;397(7):671–678. doi:10.1515/hsz-2016-0107

20. Dobin A, Davis CA, Schlesinger F, et al. STAR: ultrafast universal RNA-seq aligner. Bioinforma Oxf Engl. 2013;29(1):15–21. doi:10.1093/bioinformatics/bts635

